# Exome sequencing in families with severe mental illness identifies novel and rare variants in genes implicated in Mendelian neuropsychiatric syndromes

**DOI:** 10.1101/310821

**Authors:** Suhas Ganesh, Ahmed P Husayn, Ravi Kumar Nadella, Ravi Prabhakar More, Manasa Sheshadri, Biju Viswanath, Mahendra Rao, Sanjeev Jain, The ADBS consortium, Odity Mukherjee

## Abstract

**Introduction:** Severe Mental Illnesses (SMI), such as bipolar disorder and schizophrenia, are highly heritable, and have a complex pattern of inheritance. Genome wide association studies detect a part of the heritability, which can be attributed to common genetic variation. Examination of rare variants with Next Generation Sequencing (NGS) may add to the understanding of genetic architecture of SMIs.

**Methods:** We analyzed 32 ill subjects (with diagnosis of Bipolar Disorder, n=26; schizophrenia, n=4; schizoaffective disorder, n=1 schizophrenia like psychosis, n=1) from 8 multiplex families; and 33 healthy individuals by whole exome sequencing. Prioritized variants were selected by a 4-step filtering process, which included deleteriousness by 5 *in silico* algorithms; sharing within families, absence in the controls and rarity in South Asian sample of Exome Aggregation Consortium.

**Results:** We identified a total of 42 unique rare, non-synonymous deleterious variants in this study with an average of 5 variants per family. None of the variants were shared across families, indicating a ‘private’ mutational profile. Twenty (47.6%) of the variant harboring genes identified in this sample have been previously reported to contribute to the risk of neuropsychiatric syndromes. These include genes which are related to neurodevelopmental processes, or have been implicated in different monogenic syndromes with a severe neurodevelopmental phenotype.

**Conclusion:** NGS approaches in family based studies are useful to identify novel and rare variants in genes for complex disorders like SMI. The study further validates the phenotypic burden of rare variants in Mendelian disease genes, indicating pleiotropic effects in the etiology of severe mental illnesses.

## Introduction

Bipolar Disorder (BD) and Schizophrenia (SZ) are severe mental illness (SMI) syndromes with a median lifetime prevalence of 2.4 and 3.3 per thousand persons respectively (^Saha et al., 2005, Merikangas et al., 2011^), and an estimated heritability of 70% - 90% (^Sullivan et al., 2003, Barnett and Smoller, 2009^). Evidence from family and molecular genetic studies suggests shared, perhaps overlapping, risk factors across these syndromes (^Lin and 2007,Lichtenstein et al., 2009^). The outcomes from large-scale Genome Wide Association Studies (GWAS) exploring the Common Disease Common Variant (CDCV) hypothesis detect a proportion of the estimated genetic risk (^Cross-Disorder Group of the Psychiatric Genomics et al., 2013^). In this context, next generation sequencing (NGS) technology has enabled a deeper examination of complex disorders like SMIs by evaluating both common and rare genetic variants under the assumption of ‘oligogenic quasi-Mendelian model’ of inheritance (^Kerner, 2015^). Several recent studies in autism, SZ, BD and depression have detected rare variants using NGS in case-control or family based designs, across different genes implicated to play a key role in critical biological pathways (^Chapman et al., 2015, Kato, 2015^). Findings from such studies have shown that majority of the rare variants identified are private to a family (summarised in supplementary table1) (^Timms et al., 2013,Goes et al., 2016,Rao et al., 2017^) indicating the underlying heterogeneity in the genetic architecture of SMI. Multiplex families may provide valuable insights into the genetic correlates of these syndromes (^DeLisi, 2016^) when tested using high throughput sequencing.

A cross-nosology approach has been quite informative in identifying potential disease relevant pathways in SZ and BD (^Cristino et al., 2014, O’Dushlaine et al., 2015^). Single Nucleotide Polymorphisms (SNPs) associated with these two syndromes show a high mutual correlation, among combinations of neuropsychiatric syndromes (^Cross-Disorder Group of the Psychiatric Genomics et al., 2013^). Such overlaps have also been observed across diverse neuropsychiatric syndromes, for both common and rare genetic variations; as well as for gene expression profiles in the cerebral cortex (^Gandal et al., 2018^). These findings indicate an underlying shared molecular pathology in the patho-biology of SMIs.

As part of a longitudinal study ‘Accelerator Program for Discovery of Brain disorders using Stem Cells’ (ADBS) (^Viswanath et al., 2018^) aimed at understanding the developmental trajectories and basic biology of SMI, we describe in this study, the results of WES (Whole Exome Sequencing) in 8 multiplex pedigrees with SZ and BD phenotypes from a well characterized Indian cohort. Such studies have been predominantly conducted in large cohorts of European origin, and representation from other populations is perhaps necessary to validate earlier findings, and identify population specific signatures underlying SMIs. In the current study we aimed to identify rare, damaging, exonic variants that co-segregate with SMI in multiplex families, and examine their relevance to the disease.

## Methods

### Sample selection

The families were recruited as part of a longitudinal study ADBS, approved by the ethics committee of National Institute of Mental Health and Neurosciences, Bengaluru. The details of screening, informed consent, recruitment, and phenotyping have been previously published (^Viswanath et al., 2018^). Some of the families in the cohort have been on follow up for longer than 10 years. We have previously noted evidence of linkage in psychosis at chromosome 18p11.2 (^Mukherjee et al., 2006^), and gender-specific association to *DISC1* gene using a case control study design (^Ram Murthy et al., 2012^) in samples taken from this cohort. For the current study, eight families (A through H) with high loading of SMI (figure 1.A, supplementary figure 1) were assessed in detail. From these, 32 persons (‘cases’; 16 females) with SMI were available for blood sampling, and were subjected to whole exome sequencing. Two senior psychiatrists evaluated all patients and unaffected relatives independently. Diagnoses were made with the ICD-10 Classification for Mental and Behavioural Disorders and were verified in the longitudinal course of follow-ups. From five of the eight families, we could also sample 8 unaffected individuals who had crossed the age at risk, and defined as a ‘**family specific control**’ for the respective pedigree henceforth in this report. An independent set of 25 persons without a history of SMI were further sampled as population matched controls. Together this group constituted a total of 33 asymptomatic ‘**controls**’.

**Figure 1.**
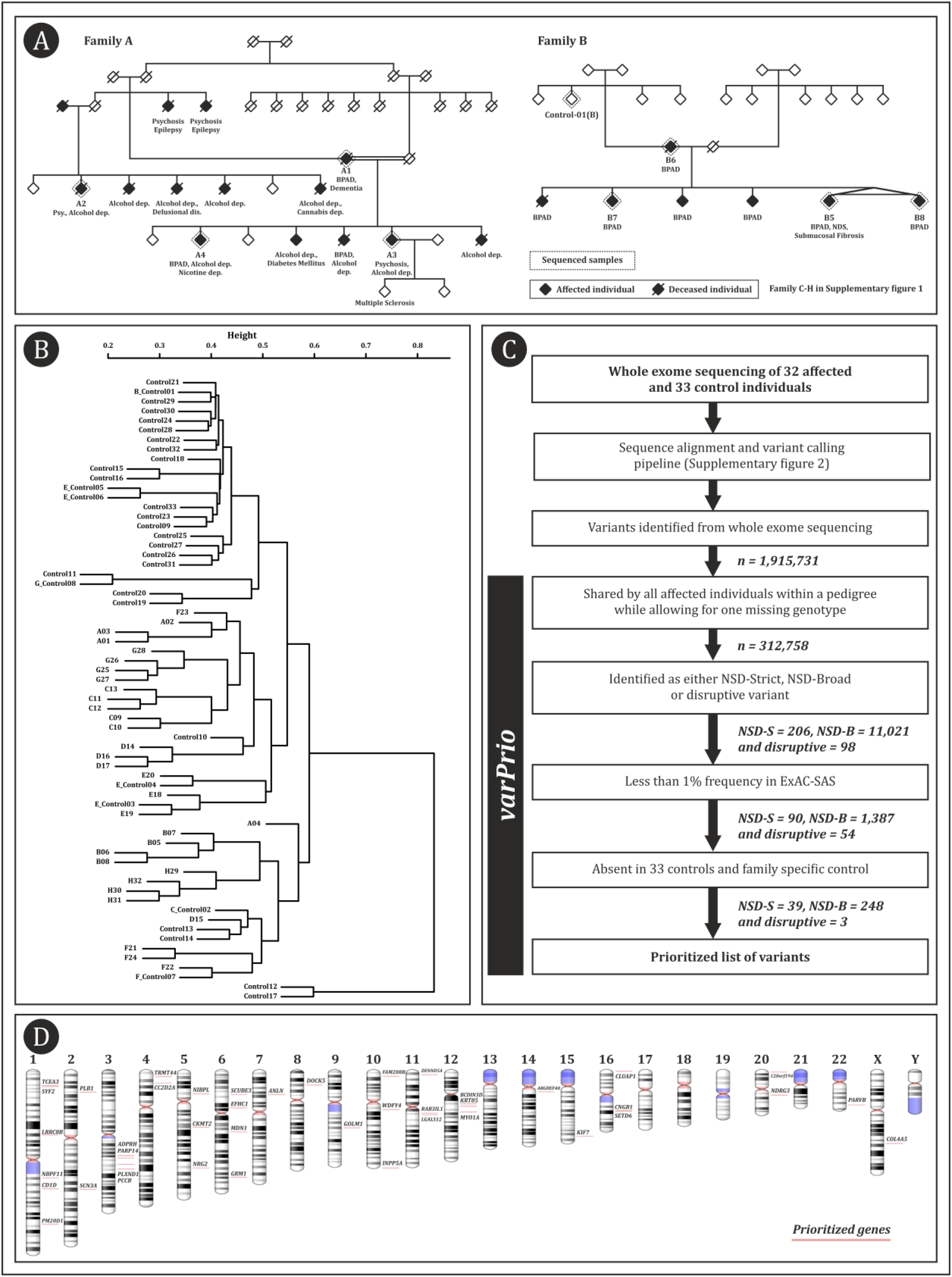
1.A Two representative pedigrees analysed with exome sequencing (A and B). 1.B Cluster dendrogram created with a distance matrix based on the degree of variant sharing between pairs of cases and controls analysed in the study 1.C ‘varPrio’ – variant prioritization pipeline with numbers indicating the reduction in the total number of variants in each prioritization step 1.D Ideogram representing the 42 genes which harboured variants prioritized by NSD-S and disruptive definition generated with NCBI genome decoration page

### Exome sequencing and analysis

Sequencing was carried out on Illumina Hiseq NGS platform with libraries prepared using Illumina exome kits. Reads were aligned with reference Human genome GRCh37 (hg19) using Burrows Wheeler Algorithm (BWA) tool (^Li and Durbin, 2009^). Variants were called from realigned BAM files using Varscan2 with the standard criteria (min coverage = 8, MAF >/= 0.25 and p</= 0.001) (^Koboldt et al., 2013^). Standard quality control protocols were employed at sequencing, alignment and variant calling (supplementary methods, supplementary figure 2). The resulting Variant Called Files (VCF) were annotated with ANNOVAR (^Wang et al., 2010^).

### Pedigree based analysis

All variant segregation analysis was performed at the level of individual pedigrees. To ascertain the degree of variance (between pedigrees) and relatedness within family structures, we performed cluster dendrogram analysis using an allele sharing matrix of the exonic variants (supplementary methods).

### Variant prioritization

Variants were prioritized if –

(a) the variant was found shared by all affected individuals within the pedigree while allowing for one missing genotype, a method shown to be useful in an earlier study of familial bipolar disorders (^Goes et al., 2016^);
(b) the variant fell into any of the following deleterious categories – **Non-Synonymous Damaging Strict** (NSD-S) set predicted to be damaging by 5 prediction algorithms - SIFT (^Kumar et al., 2009^), Polyphen-2 HDIV (^Adzhubei et al., 2010^), Mutation taster2 (^Schwarz et al., 2014^), Mutation assessor (^Reva et al., 2011^) and LRT (^Chun and Fay, 2009^) (supplementary methods); **Disruptive set** predicted to result in protein truncation (splice site, stop gain or stop loss variants) or **Non-Synonymous Damaging Broad** (NSD-B) set predicted to be damaging by one or more of the 5 prediction algorithms;
(c) the variant was absent in the 33 asymptomatic controls sequenced in this study and found rare in large population database MAF (minor allele frequency) <1% in Exome Aggregation Consortium - south Asian sample (ExAC-SAS) (^Lek et al., 2016^).

The above variant prioritization was carried out using an in-house automated pipeline ‘varPrio’. Details of the pipeline and the resulting enrichment of candidate variants is summarized in figure 1.C. To rule out any false positive calls at the final variant list, a representative set of prioritized variants (n=10) were independently confirmed by Sanger sequencing, while the remaining variants were examined for raw read depth (DDP), read depth with Phred score >20 (DP), and depth of variant supporting bases (AD).

### Functional annotation

We adapted two approaches for evaluating functional impact to the identified variants –

(a) We reviewed the literature on individual genes identified in the NSD-S and disruptive set carrying rare variants of highest priority (all 5 *in silico* predictors) for prior evidence of disease association in neuropsychiatric phenotype.
(b) For the NSD-B set carrying rare variants of plausible disease relevance (1-5 *in silico* predictors), we tested for enrichment of the aggregate list using DAVID functional annotation tool 6.8 (^Huang et al., 2009b, 2009a^). To test the enrichment on the categories of biological process, molecular function, protein domain, protein-protein interaction and tissue expression we selected the sources as - ‘ GOTERM_BP_DIRECT ‘, ‘ GOTERM_MF_DIRECT ‘, ‘INTERPRO’, ‘KEGG_PATHWAY’, and ‘UP_TISSUE’ in this *in-silico* approach. Modified Fisher’s exact test with Benjamini-Hochberg correction built-in to this algorithm was used to infer enrichment.

## Results

### Sample characteristics

Of the 32 cases sequenced in the study, 25 were diagnosed with BD, 4 with schizophrenia, 2 with schizophrenia like psychosis and one with schizo-affective disorder. They had been ill for mean (SD) duration of 23.7 (11.1) years, and the mean (SD) age at onset was 23.1 (7.9). In most of the pedigrees, there was heterogeneity in the age of onset, illness severity, global outcomes and segregation of suicidality and psychosis (in BD) with the primary phenotype. Substance use disorder was a common comorbidity, followed by hypothyroidism, seizure disorder, and dementia (supplementary table 2).

In the analysis of relatedness using the cluster dendrogram, ‘cases’ and ‘controls’ formed a single cluster possibly resulting from sharing of large number of common and/or benign variants. As expected, members from each pedigree clustered together due to relatively larger magnitude of variant sharing. (figure 1.B, supplementary methods).

### Rare deleterious variants in Mendelian genes segregate within SMI families

Family wise prioritization identified a total of 39 NSD-S, 3 disruptive and 248 NSD-B variants. The NSD-S and disruptive sets of variants (Table 1, figure 1.D) spanning 42 genes were private to individual pedigrees. Twelve of these were novel (not reported in dbSNP and other published database) and the remaining were noted in very low frequencies (<1e -07 to 7.8e-03) in ExAC-SAS.

**Table 1.**
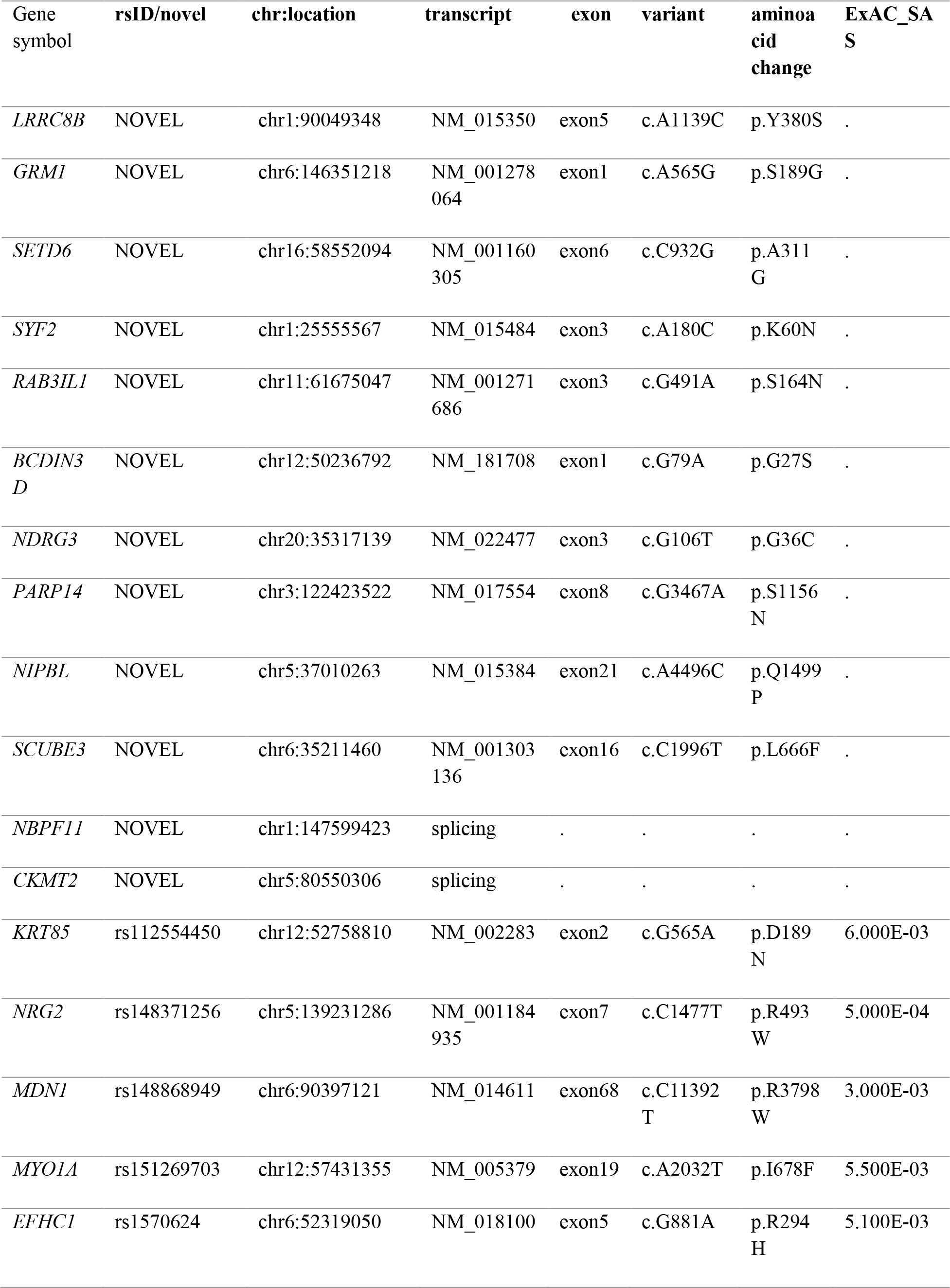
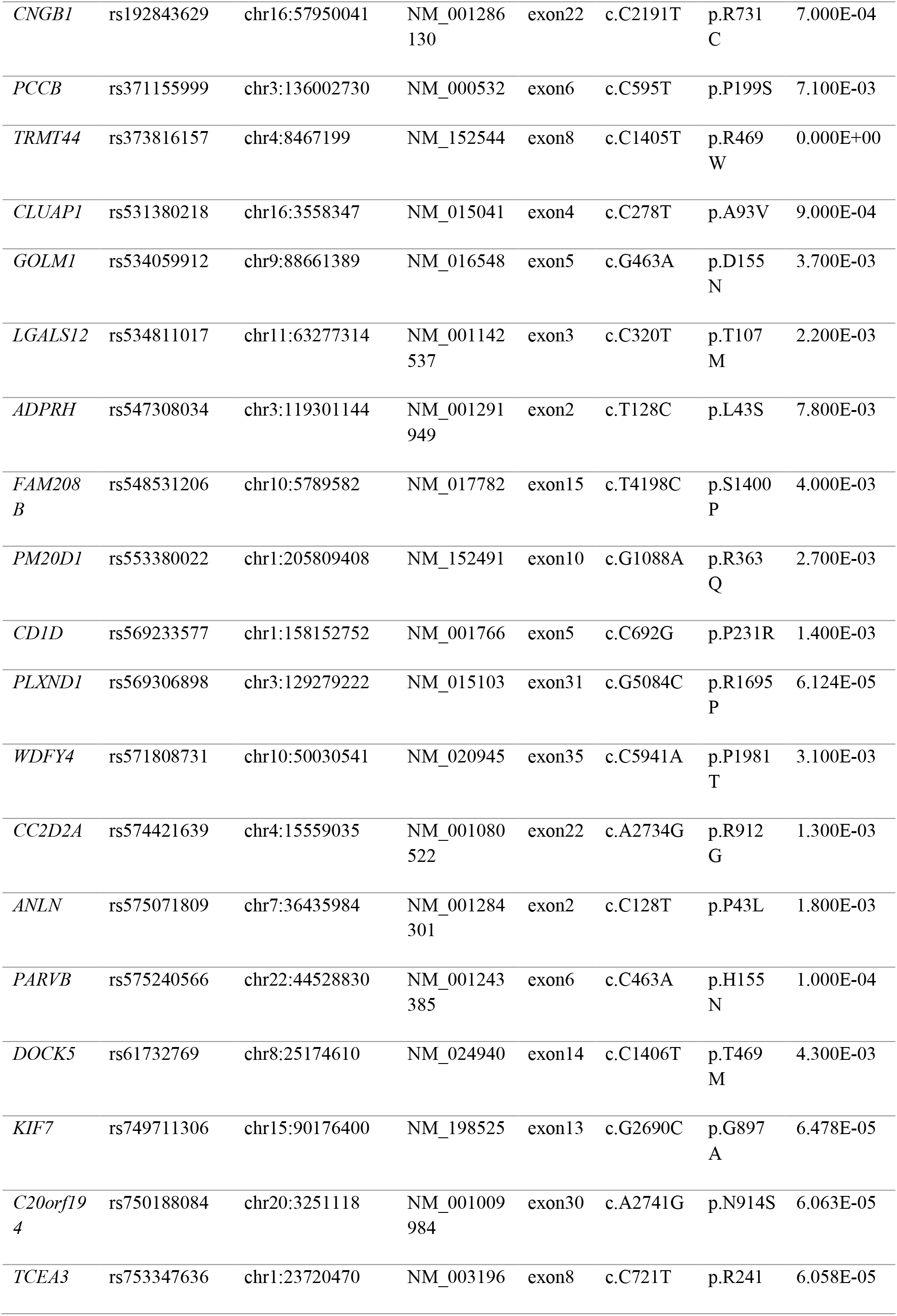
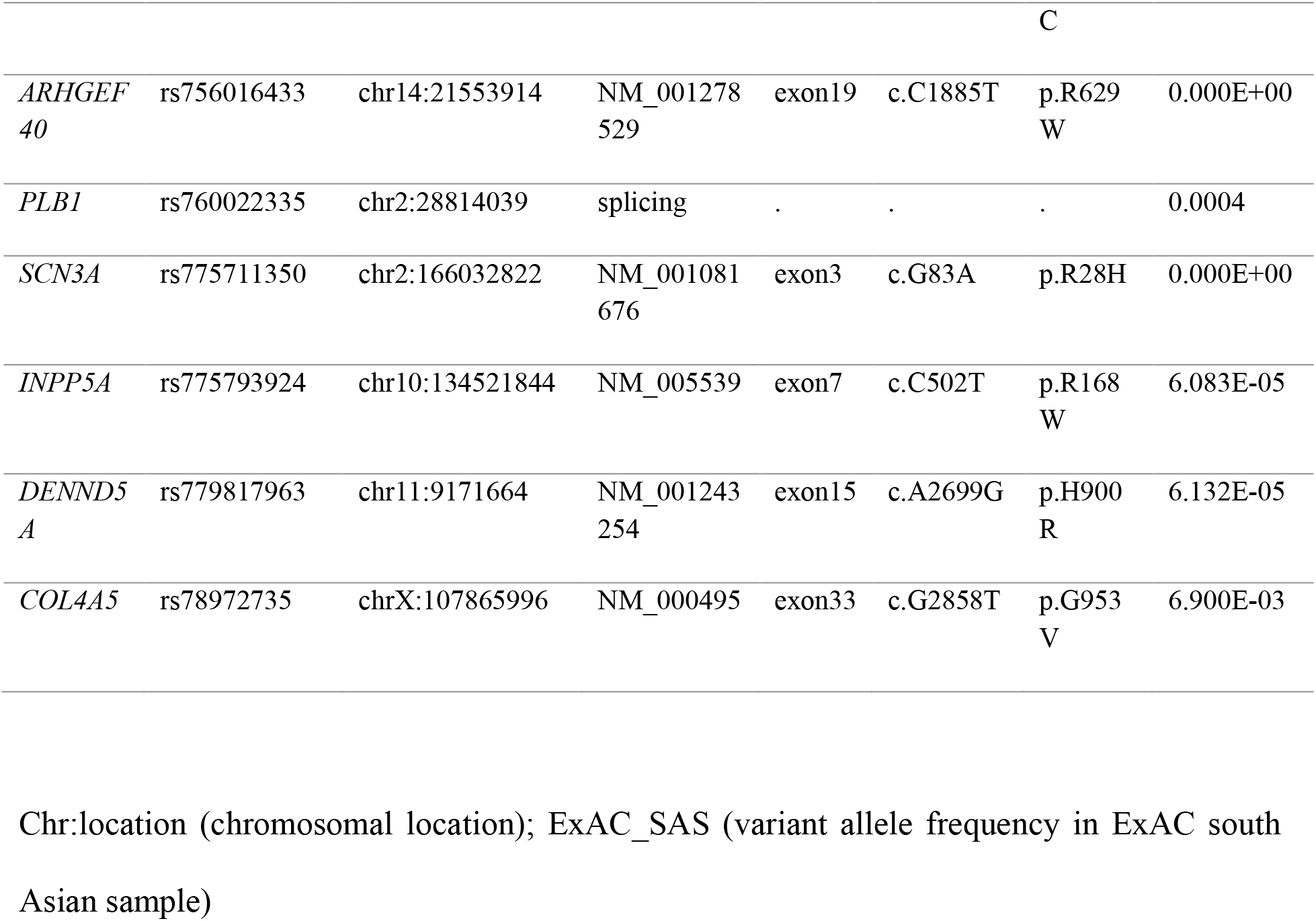
List of novel or rare, variants prioritized by NSD-S and disruptive definition

Nine of the 42 variants were found in genes which have been reported in Mendelian syndromes with early onset neurodevelopmental features such as infantile epilepsy, intellectual disability, and structural brain abnormalities. Seven of these gene-phenotype relationships were reported in Online Mendelian Inheritance in Man (OMIM) (^OMIM, 2018^) and remaining two were noted in MedGen (NCBI) and ClinVar (^Landrum et al., 2018^) databases. This was significantly higher in comparison to a background list of 1310 such genes (supplementary methods) listed in OMIM database, (P = 0.039, OR = 2.423, CI = 1.07 – 5.513, Fisher’s exact test). These variants were often observed in close proximity to reported ‘pathogenic’ mutation of the relevant Mendelian syndrome and/or in highly conserved regions (table 2.1). Two of these nine variants, one each on *GRM1* gene (chr6:146351218, GRCh37), and *NIPBL* gene (chr5:37010263, GRCh37) were novel. Pathogenic mutations on *GRM1* gene, coding for metabotropic glutamate receptor 1 (mGluR1) result in autosomal dominant (type 44) (OMIM:617691) and recessive (type 13) (OMIM:614831) forms of Spinocerebellar Ataxia both of which are characterized by early age of onset and associated intellectual disability. Missense variants have been identified spanning the entire exome of this gene in persons and families with schizophrenia and other neuropsychiatric syndromes (^Ayoub et al., 2012^). Mutations in *NIPBL* gene, coding for Cohesin Loading Factor involved cortical neuronal migration (^van den Berg et al., 2017^), cause Cornelia de’Lange syndrome 1. The novel missense variant identified in the pedigree G (chr5:37010263), segregating with bipolar disorder would result in substitution of polar amino acid glutamine by a hydrophobic amino acid proline. A non-sense mutation at the same codon (rs797045760) is reported to be pathogenic of Cornelia de’ Lange syndrome 1 (ClinVarSCV000248215.1).

**Table 2.**
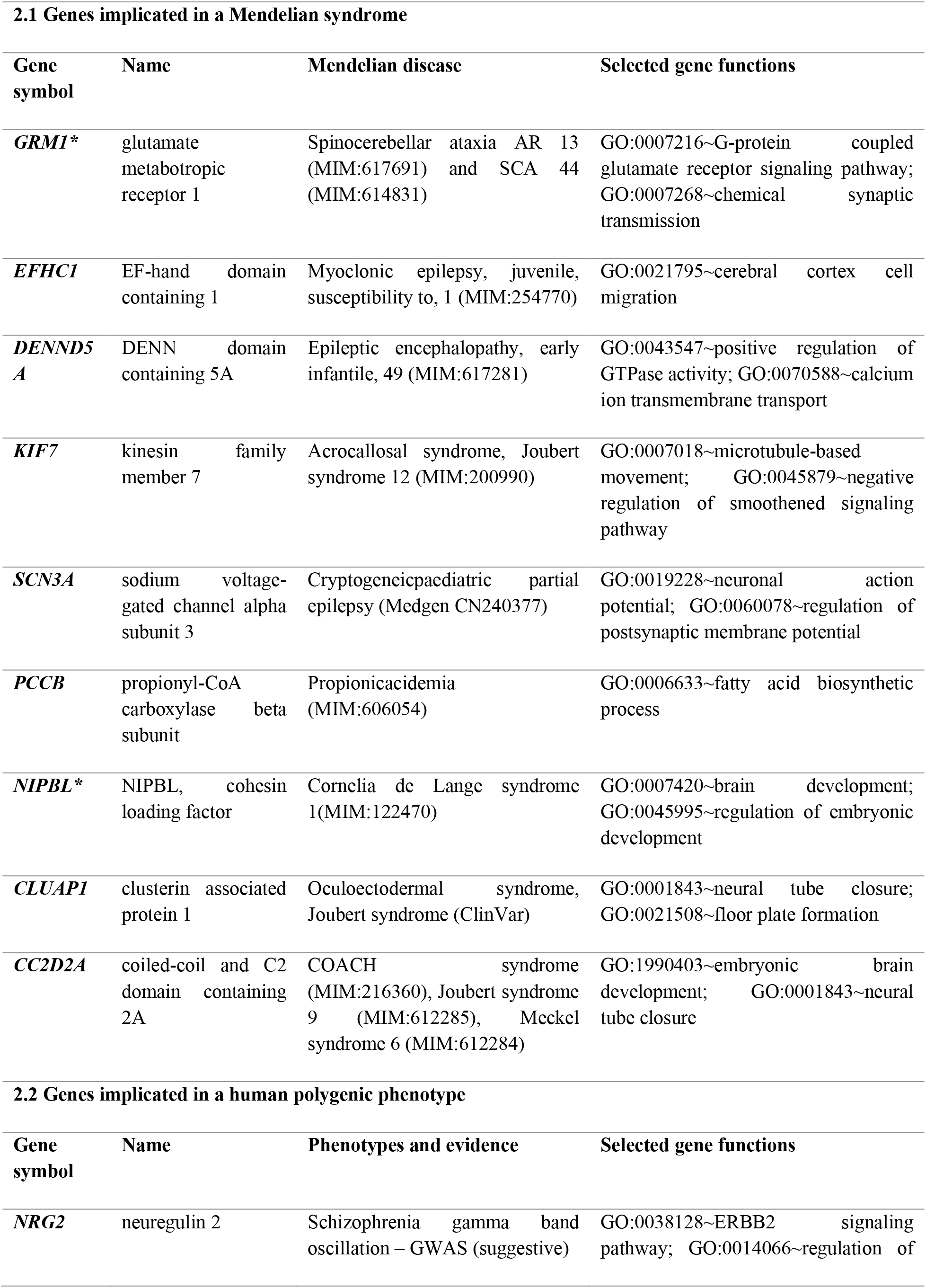
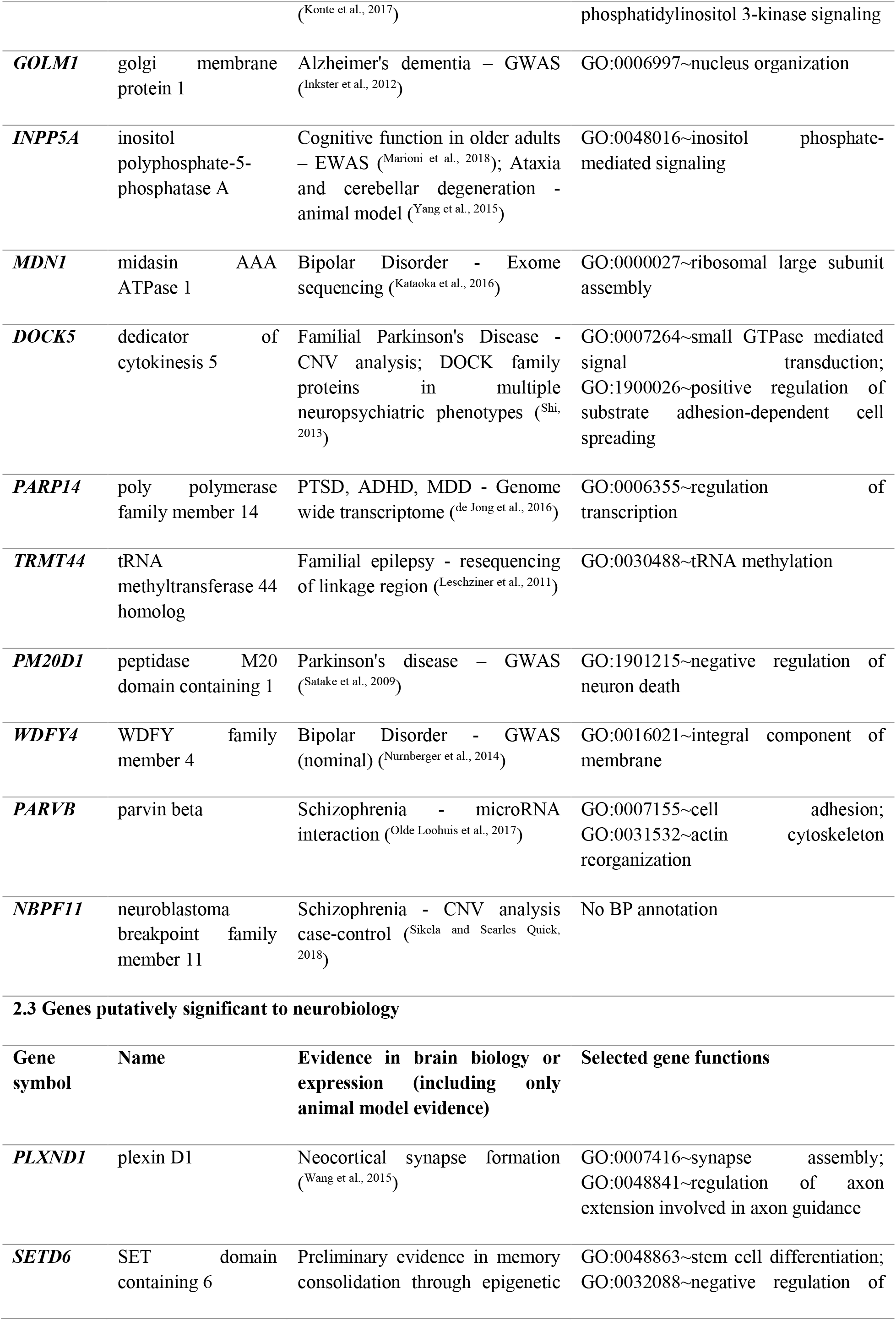
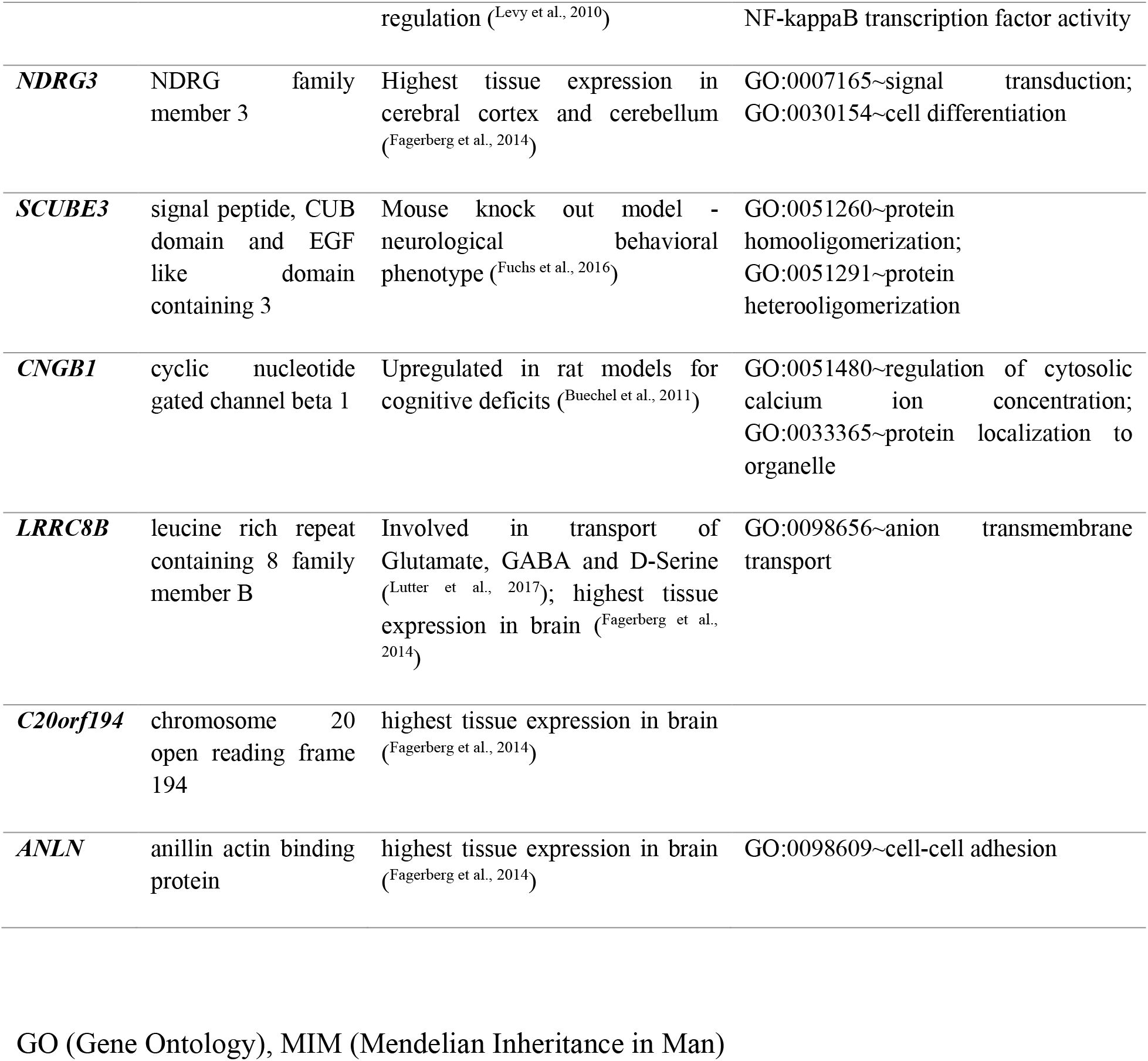
Disease relevance of the genes harbouring prioritized variants

Ten other genes that harboured prioritized variants, have been implicated in neuropsychiatric syndromes. We identified a variant (rs148371256) in *NRG2* gene (Neuregulin 2) which was earlier reported to be associated with gamma band oscillations in schizophrenia with suggestive genome wide significance (^Konte et al., 2017^) and the encoded protein neuregulin-2 has been shown to be critical for the formation and maturation of GABAergic synapses (^Lee et al., 2015^) and its ablation results in dopamine dysregulation (^Yan et al., 2017^). Another novel variant (chr3:122423522, GRCH37) was identified in *PARP14* gene (Poly ADP ribose polymerase 14), and the gene has been implicated in Post-Traumatic Stress Disorder (PTSD), Major Depression (MDD) and Attention Deficit Hyper Kinetic Disorder (ADHD) (^de Jong et al., 2016^). We also noted a variant (rs534059912) in *GOLM1* gene (Golgi membrane protein 1), which was earlier reported in sporadic Alzheimer’s dementia (AD) to influence the pre-frontal cortical volume (^Inkster et al., 2012^). A list of these 10 genes, evidence for disease association and gene ontology descriptions are presented in table 2.2.

Of the remaining genes, there were several with a plausible role in the biology of SMI, but not thus far implicated in any disease phenotype. These genes, with the ontology descriptions and plausible biological implications are provided in table 2.3.

### Enrichment of coding variants with plausible functional role in SMI

NSD-B set consisted of 248 variants, of which except for rs570064523 on *PCSK1* gene identified in cases from two families (G and H), no other overlap at the level of family was noted for the remaining 247 variants (supplementary table 3). This set of genes was subjected to enrichment analysis to evaluate biological implication. In the ‘protein domains’ category tested using Interpro database as the source, the term ‘Epidermal Growth Factor like domain’ showed a nominally significant enrichment with p = 0.0013, Benjamini Hochberg False discovery rate corrected p = 0.073. Twelve genes, which enriched for this domain included *NRG2 and SCUBE3* genes that were also categorised in the NSD-S set along with *NOTCH1, JAG1* and *WIF1* genes which form critical nodes in the Notch signalling pathway implicated in neurodevelopment(supplementary table 4) (^Lasky and Wu, 2005^). There was no statistically significant enrichment in any of the remaining categories tested with this in-silico approach.

## Discussion

The results of our study highlight the usefulness of WES in multiplex families with SMIs, to identify rare and novel variants that may contribute to susceptibility to disease. Many of these variants prioritized by NSD-S, and presumed to be disruptive, map to genes which are previously reported in GWAS, candidate gene association, post-mortem expression, or animal model studies of SMIs. In addition, consistent with the WES approach, we identify variants in genes hitherto not reported in the context of a SMI, but could potentially contribute to disease biology.

The segregation of rare and deleterious variants in Mendelian disease genes with a neuropsychiatric phenotype is in keeping with some recent observations. Studies have shown that heterozygous carriers of Mendelian disease mutations are at increased risk for specific common diseases. While Mendelian forms of common, complex traits such as Alzheimer’s disease, hypertension, hypercholesterolemia, hypertriglyceridemia have long been attributed to rare causal variants in single genes, population based GWAS studies in these traits have often implicated genes that also cause single gene disorders (^Lupski et al., 2011^). More recently, using electronic health record data, disease relevant phenotypic burden of rare variants in Mendelian genes, thus far not characterized as ‘ pathogenic’ has been demonstrated across diverse phenotypes (^Bastarache et al., 2018^).

We explored the clinical significance of nine variants in Mendelian genes in ClinVar database, a publicly available archive of human phenotype - variation relationships (^Landrum et al., 2018^). None of these variants were annotated as ‘pathogenic’ or ‘likely pathogenic’ in the database for the corresponding Mendelian phenotype. As a corollary, none of the families had any identified or suspected case of a severe neuro-developmental syndrome. However, the predicted deleteriousness by *in silico* algorithms, a very low prevalence in the population, physical proximity to known pathogenic mutations, and the reported physiological gene function suggest a plausible role for these variants in the aetiology of SMIs. The impact of these variants in cellular and/or animal models needs to be carried out to validate these observations and to establish their role in SMI. Interestingly, an earlier WES study in families with BD, also reported variants in genes of monogenic syndromes - Holoprosencephaly and Progressive Myoclonic Epilepsy (^Rao et al., 2017^).

We detected rare variants in ten additional genes that have been noted in earlier studies, to contribute to the risk for polygenic syndromes such as SZ, BD, Autism, MDD, ADHD, PTSD, AD and Parkinson’s Disease. This finding is congruent with the evolving concept of shared molecular neuro-pathology across SMIs (^Gandal et al., 2018^). These, along with other identified genes known to be involved in neurodevelopmental processes (e.g. *PLXND1*) or known to have manifold higher brain expression (e.g. *ANLN, LRRC8B)* are potential targets to be examined in future studies of SMI. Lastly, of the twelve genes encoding a highly conserved Epidermal Growth Factor like domains and showing nominally significant enrichment to this domain, many encode for proteins that play critical role during embryogenesis and neurodevelopment (^Lasky and Wu, 2005^).

Certain limitations are to be considered while interpreting the results of this study. The relatively small control set sequenced in our study precluded statistical association testing at the level of a variant or a gene. As an alternative, we considered the minor allele frequency of the variant in ExAC South Asian Sample in the prioritization approach, and many were noted to be extremely rare. This could have resulted in few false positive discoveries, and at the same time missed out few true disease associations. Secondly, although we sampled nearly equal number of affected persons from each family, the relationships within pedigrees were not uniform, potentially adding heterogeneity to the number of identified variants. Thus, we prioritized variants with complete sharing allowing for one missing genotype. Third, like the previous studies of WES in SZ and BD, we have relied on *in silico* predictions to infer the deleteriousness of a variant. Lastly, inherent to the prioritization criteria of rarity, deleteriousness and segregation the NSD-S and disruptive variant set presented above would explain only a part of an individual’s liability to disease. The results of this analysis represent the shared familial risk for SMI, private to each pedigree, determined by variants of possible major/moderate effect.

Using WES data in multiplex families with SMIs, we find evidence that suggests intersections in the molecular pathways leading to the expression of polygenic SMIs and Mendelian neuropsychiatric syndromes. The patient derived neural stem cell lines being developed as part of the program (^Viswanath et al., 2018^) will be useful to explore the functional significance of the identified variants accounting for ‘modifier genetic background’ (^Riordan and Nadeau, 2017^), and to characterize mechanisms that underlie the observed genotype-phenotype correlates.

## Conclusions

NGS approaches in a family based study design are useful to identify novel, and rare variants in genes potentially relevant to complex disorders like SMIs. The study further provides an independent validation for the phenotypic burden of rare deleterious variants in Mendelian disease genes that segregate privately in multiplex pedigrees with SZ and BD. Our findings support the role of heterogeneity and pleiotropy in the genetic architecture of SMIs encompassing a spectrum of neurodevelopmental and degenerative phenotypes.

## Acknowledgements

The authors are grateful to all the patients, their family members and healthy volunteers who participated in the study. Financial support for the study was provided by Department of Biotechnology funded grants - BT/01/CEIB/11/VI/11/2012, entitled, “Targeted generation and interrogation of cellular models and networks in neuro-psychiatric disorders using candidate genes” and BT/PR17316/MED/31/326/2015 entitled, “Accelerating program for discovery in brain disorders using stem cells” (ADBS), Pratiksha Trust and The Institute of Stem Cells and Regenerative Medicine (InStem), Bengaluru, India.

The authors would like to thank the sequencing core facility at the Institute of Genomics and Integrative Biology (IGIB), Delhi (Dr. Faruq Mohammed) and the National Centre for Biological Sciences (NCBS), Bengaluru (Dr. Awadhesh Pandit) for sample processing and WES data generation.

The authors would like to thank Manasa K P, Anand Ganapathy Subramaniam, Soham Deepak Jagtap, Geetanjali Murari, Srividya Shetty, Surya Prakash M from NIMHANS & Vidhya Varadharajan, Priyanka Bhatia, Shubhra Acharya and Batul Yusuf from InStem for their technical support during initial raw data curation.

The authors would like to thank all investigators of ADBS consortia for providing valuable inputs to the manuscript and having final approval of the manuscript. SG is currently affiliated with and supported by Schizophrenia Neuropharmacology Research Group at Yale university.

